# Canopy traits reveal climate-associated genetic differentiation in a California oak common garden

**DOI:** 10.64898/2026.10.20.760433

**Authors:** Alexander R. B. Goetz, Derek J. N. Young, Kara K. Sanghera, Jessica W. Wright, Victoria L. Sork

## Abstract

**Background and Aims:** Phenotypic variation in plant populations reflecting historical climate adaptation may affect performance under future climates. In long-lived trees, height and stem diameter are commonly used to assess growth, but these measures may not fully capture complementary structural and functional dimensions of performance, such as canopy development and vigor. We investigated whether canopy traits reveal climate-associated genetic differentiation in *Quercus lobata*.

**Methods:** Using drone-based remote sensing of 1,395 12-year-old *Quercus lobata* trees from 601 families in a common garden, we quantified canopy volume and greenness, using the Normalized Difference Vegetation Index (NDVI). Previous analyses indicated that trees originating from warmer climates grew more in common gardens than those from cooler climates. To assess whether canopy traits showed similar patterns, we tested for family-level variation in each trait and associations with climate of origin.

**Key Results:** Canopy volume and greenness showed significant genetic variation among families and significant associations with climate of origin. Trees from hotter climates had greater canopy volume and greenness than trees from cooler climates.

**Conclusions:** These covarying traits provide novel evidence of climate-associated genetic differentiation in tree performance, consistent with historical climate selection. Trees originating from climates with hot summers, moderate winters, and low precipitation performed best in the garden, suggesting that they will perform better than trees adapted to cool, wet conditions under future climate change.

## 1. INTRODUCTION

The fate of any organism is determined by the extent to which it is adapted to the environment where it lives (Williams 1966, Mátyás 1996, Kawecki and Ebert 2004, Sork 2018). Rapid shifts in climate may cause long-lived sessile organisms to experience different environmental conditions than those to which they are adapted (Aitken et al. 2008, Wang et al. 2010, Franks et al. 2014, Mahony et al. 2020, Erlichman et al. 2024). If populations are unable to evolve quickly enough to the new environment, they will become less fit to their local environments and could become extinct. Therefore, the conditions of a genotype’s origin are likely to shape its performance in an altered environment (Rehfeldt et al. 1999, 2002, Schwinning et al. 2022).

Common garden studies, where multiple genotypes are planted in the same environment to compare their phenotypes, provide a powerful tool for assessing how climate is associated with tree performance (Schwinning et al. 2022) and the genetic basis of phenotypic traits affecting that performance (e.g., Azuma et al. 2023, Capblancq et al. 2023, Grubinger et al. 2023). Physical size of trees is an informative surrogate for fitness because it predicts survival and fecundity (e.g., Younginger et al. 2017). Height and stem diameter, common easy-to-measure metrics for assessing tree size, are known to scale with total biomass (Chave et al. 2014). However, repeated measurements from the ground can be labor-intensive, especially for large sample sizes (Huxman et al. 2022), and ground measurements of canopy volume or architecture can be especially difficult. Therefore, it is productive to consider other methods of estimating tree performance in common garden studies to predict relationships between climate and performance.

In contrast to measurements on the ground, remote sensing using uncrewed aerial vehicles (UAVs; referred to throughout as “drones”) provides a high throughput means of phenotyping trees grown under experimental conditions. Drones and crewed aircraft allow us to evaluate structural and functional traits that may be under selection (Baulies and Pons 1995). Most immediately, we can quantify the volume of the entire tree canopy, which scales closely with total aboveground biomass (Zimble et al. 2003, Wang et al. 2008, Bouvier et al. 2015), as well as characterize tree architecture. In angiosperms, some individuals grow tall while others produce many horizontal branches, and this variation in architecture may confound allometric size relationships (Valladares and Niinemets 2007, Segura et al. 2009, Costes and Gion 2015). Thus, aerial imaging may provide more detailed information than stem diameter and height.

Aerial images of tree canopies capture other dimensions of functional phenotypes such as high photosynthesis, good or bad vegetative condition, plant stress, and other leaf conditions (Seidel et al. 2011, Laurans et al. 2024). Specifically, greenness of leaves (measured as the Normalized Difference Vegetation Index; NDVI) is a complex phenotype that is mechanistically linked to leaf health and physiological processes related to carbon assimilation (Knipling 1970, Sellers 1985, Carter 1993, Peñuelas and Filella 1998, Gao 2000, Wang et al. 2004). Here, we primarily discuss greenness as a proxy for canopy health at the time of measurement. In other tree species, greenness has been shown to be genetically differentiated among individuals (e.g., Corbin et al. 2025) and scale closely with annual growth (Wang et al. 2004). Volume and greenness are both expected to be highly polygenic traits (Cornelius 1994, McKown et al. 2014). Thus, measuring both volume and spectral signatures at the individual canopy level in a well-replicated common garden study allows us to test whether these metrics reveal biologically meaningful genetic and climate-associated differences in structural and functional aspects of tree performance and vigor.

*Quercus lobata* (valley oak), an ecosystem-foundational species endemic to California, is distributed across a range of climate environments (Pavlik et al. 1990) that are likely to incorporate large intraspecific variation in climate adaptation. Numerous climate-associated morphological and functional traits measured in the same common gardens show significant genetic differentiation among facilities associated with climate of origin, including leaf thickness, leaf area, trichome density, and water use efficiency (MacDonald 2017, Ochoa 2024, Badillo 2026). We have also reported genetic differences in leaf phenology associated with climate of origin (Wright et al. 2021, Goetz et al. 2026b). Previous work from these gardens has shown evidence of heritable growth variation based on stem diameter and height differences among families (Browne et al. 2019, Goetz et al. 2026a), and report evidence of maladaptation to current climates (Browne et al. 2019, Goetz et al. 2026a). Thus, both the distribution of the species and the prior research make the *Q. lobata* study system ideal to investigate drone-based measures of tree performance.

In this study, we apply high-throughput drone-based phenotyping to a twelve-year, range-wide *Q. lobata* common garden experiment (described in Delfino Mix et al. 2015). By quantifying structural and functional dimensions of tree performance using drone-based phenotyping methods, we can test the effects of genetics and climate of origin on canopy-scale phenotypes to better understand how tree species will respond to climate change. We asked whether high-throughput canopy phenotyping can reveal climate-associated genetic variation in tree performance. Using photogrammetry and multispectral imaging, we quantified canopy volume and greenness in 1,088 12-year-old Q*. lobata* trees representing 557 half-sib families in a range-wide common garden. We predicted that the complementary high-throughput traits of canopy volume and greenness would be genetically differentiated among families and that variation in both traits would be associated with climate of origin, consistent with climate-associated genetic differentiation among source populations. Together, these high-throughput data provide additional phenotypes to examine tree performance in terms of canopy volume, a measure of tree size, and greenness, a measure of canopy vigor, and provide a scalable approach for characterizing population responses to changing climates. Such results can be used to inform management strategies for oak populations that are predicted to experience warmer climates.

## 2. MATERIALS AND METHODS

### 2.1. Study species

*Quercus lobata* Née (valley oak) is a long-lived, winter-deciduous tree oak endemic to woodlands, savannas, and riparian areas of the California Floristic Province (Pavlik et al. 1991). It is considered a foundational species in the California oak savanna and riparian ecosystems because it supports extraordinary biodiversity (Allen-Diaz et al. 1999).The species is found across wide abiotic gradients (0-1700m above sea level; Pavlik et al. 1991). *Quercus lobata* has shown widespread genetic differentiation across differing climates (e.g., MacDonald 2017, Wright et al. 2021, Ochoa 2024, Ochoa et al. *in press*), but this local adaptation has lagged behind historical warming since the Last Glacial Maximum (Browne et al. 2019). It is of conservation concern due to land use change (Whipple et al. 2011), recruitment limitation (Tyler et al. 2006, Zavaleta et al. 2007), and climate change (Sork et al. 2010, Goetz et al. 2026a). Mature *Q. lobata* trees reach heights of 12-30 m, typically with high degrees of lateral branching (Pavlik et al. 1991).

### 2.2. Common garden

This study was conducted in a *Q. lobata* common garden experiment established by Sork and Wright, now in its 15^th^ year (see Delfino Mix et al. 2015 for detailed methods and experimental design). The garden used in this study (USDA Institute of Forest Genetics, Placerville, CA) is 822m elevation above sea level and has an intermediate climate relative to the rest of the *Q. lobata* range (Delfino Mix et al. 2015).

In 2015, two-year-old half-sib seedlings of 674 adult trees across the species range were planted into the common garden plot (Figure 1). Acorns were collected in 2012 from open-pollinated adult trees. Multiple acorns from each tree were germinated, thus the trees in the garden comprise half-sib families of the adult trees. Tree growth and survival have been measured annually by field crews from 2013 to present. As planned, trees were deliberately thinned in 2021 to prevent future competition from neighbors. At the time of data collection in 2024, 1,668 living trees from 622 families were present in the garden, and each family had 1-3 living progeny.

**Figure 1.**
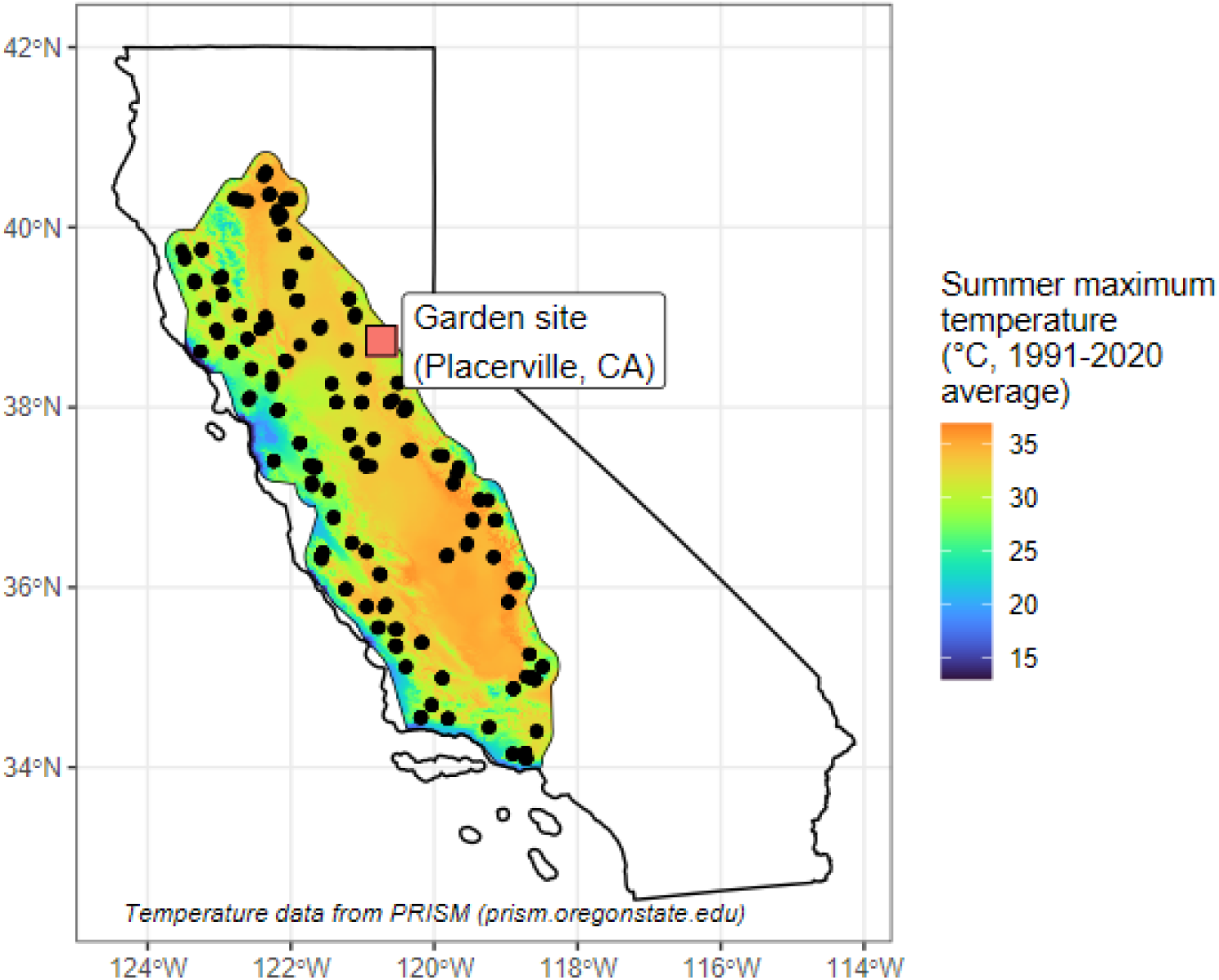
Location of common garden site (red square) and mid-points of 95 locations (black dots) where acorns were collected from 573 family lineages. Color represents average summer maximum temperature 1991-2020 (PRISM; prism.oregonstate.edu). Temperature is shown within the current range of *Quercus lobata*, extrapolated from occupancy data.

**Figure 2.**
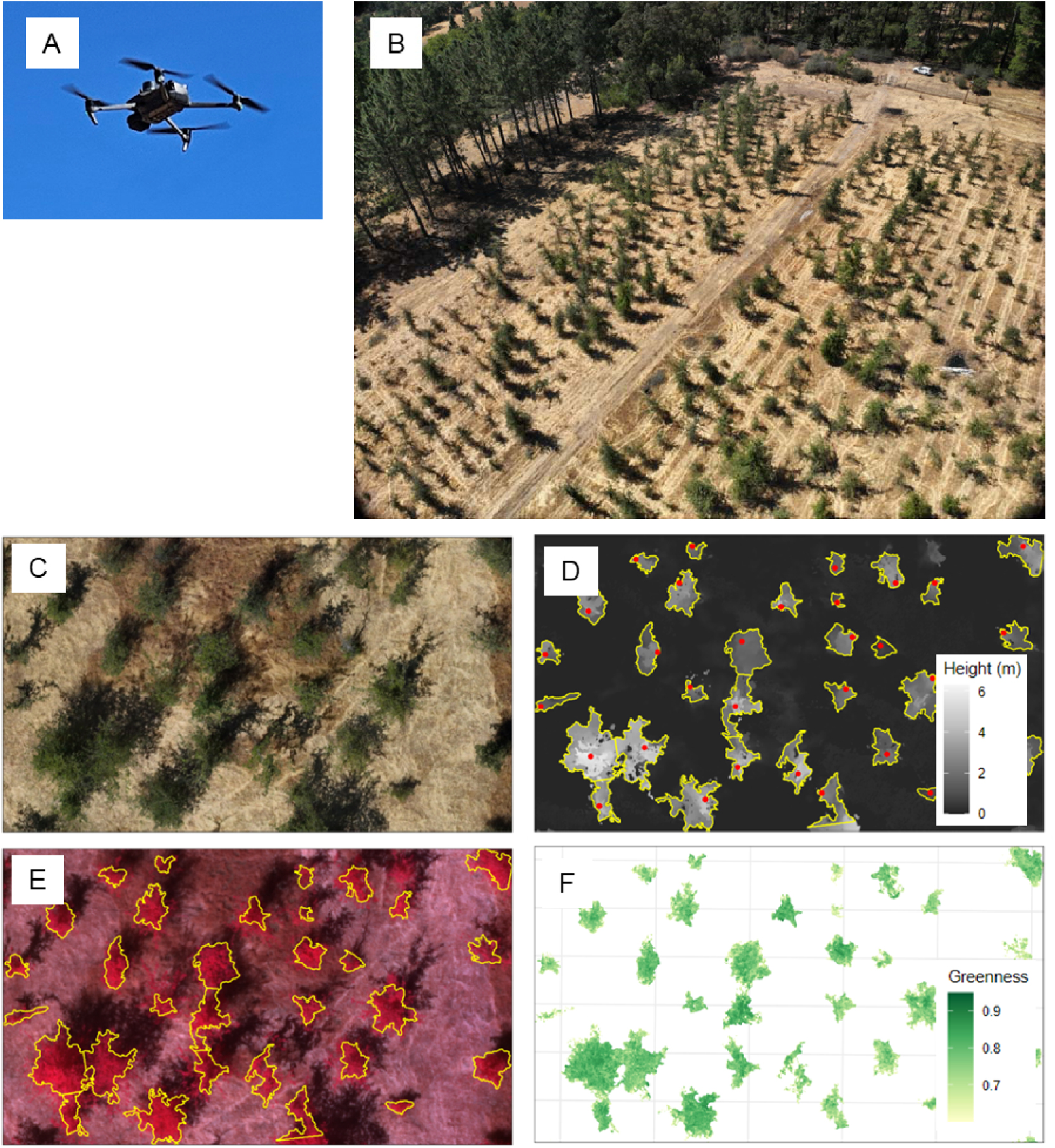
Overview of methods to measure size and greenness of 12-year-old *Quercus lobata* trees growing in a common garden. (**A**) Drone (DJI Mavic 3 Multispectral) in flight during data collection (photo: A. Goetz). (**B**) Aerial view of a section of the common garden, taken during flight. C-F: Remotely sensed images. (**C**) True-color raster image of a subset of study trees. (**D**) Canopy height model raster image used to determine tree height, canopy extent, and canopy volume. Red points are treetops (local maxima in CHM, used to segment CHM into canopies). Yellow outlines are canopy boundaries calculated from the CHM using the algorithm of Silva et al. (2016). (**E**) False-color multispectral orthomosaic showing near-infrared, red, and green bands. Yellow outlines are canopy boundaries; greenness was only calculated on pixels within tree canopies. (**F)** Derived raster of canopy greenness (normalized differentiation vegetation index) by pixel. Pixels with greenness < 0.6 were dropped from further analyses because they were unlikely to represent living vegetation (Gao 2000). Greenness values were then averaged by canopy for further analyses.

### 2.3. Climates of origin

To characterize the climate of origin affected phenotypes of trees in the common garden, we acquired historical temperature and precipitation data at the origin coordinates from the Basin Characterization Model (Flint et al. 2013). The Basin Characterization Model uses a large set of elevation and hydrological data to downscale climate interpolations to a 270m raster grid, making it a highly accurate representation of climate across the state (Flint et al. 2013). We used average values from the earliest period (1951-1980) available in this high-accuracy, high-resolution dataset to represent climate of the maternal trees. We used these historical climate data because all maternal trees were estimated to be at least 50 years old (Sork and Wright, unpublished data), so 1951-1980 data is a closer approximation of the climatic conditions in which the trees would have germinated (e.g., Sáenz-Romero et al. 2015, 2019, Horváth and Mátyás 2016, Kooyers et al. 2019). Because historical climates were more stable than modern climates (LaDochy and Witiw 2023) and 1951-1980 climate data were closely correlated with climate data from earlier time periods (Browne et al. 2019, Goetz et al. 2026a), the specific choice of historical time period was less important than capturing the difference between historical and current temperature. To compare recent-past maternal temperature data with current average temperatures at the common garden site, we obtained 2014-2024 averages of temperature and precipitation data at the garden’s point location from the PRISM database (prism.oregonstate.edu) at 800m pixel resolution with grid cell interpolation. The study year (2024) had high maximum summer temperatures relative to the rest of the experimental period (Goetz et al. 2026a), but annual precipitation was higher than average (Figure S1).

We used the following variables: average of monthly temperature maxima in the summer months (June-August), referred to as “T_max_”; average temperature in the coldest quarter (bioclimatic variable 11; Fick and Hijmans 2017), referred to as “T_min_”; and cumulative yearly precipitation. We chose these variables because they reflect the main climatic determinants of *Q. lobata* distribution (Kueppers et al. 2005, Sork et al. 2010, McLaughlin and Zavaleta 2012, Gugger et al. 2013). Temperatures are also projected to increase and precipitation is projected to decrease in the future throughout the species range (Wang et al. 2017), so these variables are relevant for examining the effects of climate change on performance.

### 2.4. Data collection

Remotely sensed data were collected in August 2024. We used a Mavic 3 Multispectral drone (DJI, Shenzhen, China), set to collect imagery with a 90/90 front/side overlap (%) at 60 m above ground level, collecting a red-green-blue (RGB) and multispectral (MS; consisting of green, red, red-edge, and near-infrared (NIR) bands) image at each photo point (with photo points determined based on overlap computed using the RGB camera’s field of view). Images of a spectral calibration panel with known reflectance values (T3-R50; MAPIR, Inc., San Diego, California, USA) were collected at the start and end of the data acquisition mission.

In September 2024, we measured height on all trees in the common garden using telescoping height poles (three trees above 7.6m tall were instead measured with a Forestry Pro II laser hypsometer; Nikon Corporation, Tokyo, Japan). For trees at least 1.5m tall, we measured diameter at breast height (DBH; 1.4m above ground) of the dominant stem. We also identified locations of volunteer plants (i.e., plants not deliberately planted for the study, either *Q. lobata* or other species). In October-December 2024, we recorded locations of all trees using a Geode GNS3 handheld GPS unit (Juniper Systems, Inc., Logan, Utah, USA). The accuracy of the GPS relative to the geoid in low to no canopy cover was ±0.30m. We took each GPS point at a south-facing part of the stem. For consistency in measurement, GPS points were only taken on sunny or mostly sunny days.

### 2.5. Data processing

Imagery was processed in Metashape v2.2 (Agisoft LLC: St. Petersburg, Russia). All images (RGB and MS) were aligned together with quality setting "high". Multispectral images were radiometrically calibrated against the known reflectance values of the calibration panel, with pre-and post-mission calibration values averaged and data from the drone’s downwelling light sensor used to adjust for within-mission fluctuations in illumination. Then, a dense point cloud, digital surface model (DSM; height raster of ground and vegetation surface features), and digital terrain model (DTM; height raster of terrain without surface features) were created using only the RGB images with quality setting "high" and filtering "moderate". The resulting DSM was used as the 3D surface for creating the stitched orthomosaic from the RGB images and (separately) the MS images. We used computational resources from Jetstream2 at Indiana University (Hancock et al. 2021, Boerner et al. 2023) for imagery processing.

To detect treetops, we computed a canopy height model (CHM; height raster of vegetation normalized by terrain) by subtracting the DTM from the DSM at the native resolution, then resampled (coarsened) to 0.1 m resolution. Using the resulting CHM, we detected treetops using a fixed 2-m radius local maximum filter with a minimum height threshold of 0.5 m. Using the resulting treetops, we delineated crown extent using the algorithm of Silva et al. (2016), implemented in function silva2016() in R package lidR (Roussel and Auty 2020), with no restriction on crown diameter relative to tree height and minimum crown height set to 25% of treetop height. We applied a smooth (default parameters of function smooth() in package “smoothr” version 1.1.0; Strimas-Mackey 2025) to the resulting polygons and then simplified the smoothed polygons using a tolerance of 0.01 m.

To ground-truth the automated treetop detection, we manually checked for errors by visually inspecting the canopy polygons and treetop points in relation to the RGB orthomosaic and by comparing canopy size and treetop height with the ground-truth field data. We spatially joined the treetop points, canopy polygons, and RGB orthomosaic to the ground-truthed GPS points using functions from the “sf” package (version 1.0-22; Pebesma 2018, Pebesma and Bivand 2023). We first removed all points/polygons outside the garden plot or marked as volunteer plants during field data collection. We then ground-truthed and filtered the remaining verified study trees.

Among verified study trees, the most common sources of error were (a) a single tree incorrectly marked as multiple (due to structural irregularities in the canopy) or (b) multiple trees incorrectly marked as one (due to multiple canopies growing close together). We remedied situation (a) by combining (function union() in package “sf” version 1.0-22; Pebesma 2018, Pebesma and Bivand 2023) all polygons representing a single tree canopy and treating the highest treetop point as the new “overall” treetop. We remedied situation (b) by re-running the treetop detection algorithm on the incorrectly classified treetops with less restrictive parameters (thus generating more “candidate” treetops) and retaining the *n* highest treetops, where *n* = number of distinct trees known to occur within a polygon based on the ground-truth data. Using these new treetop points, we re-ran the canopy segmentation algorithm on the CHM with the same parameters as before, including smoothing and simplifying. We then visually re-checked for concordance with the known locations of trees.

We excluded a total of 502 trees in the garden from analysis because we were unable to confidently determine their crown extents and sizes. First, 249 small trees were automatically excluded because the treetop detection algorithm and crown segmentation algorithm did not detect them. We excluded 53 additional trees with unresolved crowns (i.e., the re-run treetop detection algorithm still resulted in 1 treetop, typically because a taller tree’s canopy obscured a shorter tree’s canopy). We also removed 57 trees with higher drone-estimated heights than ground-measured heights, which indicated incorrect measurement by the drone. Finally, 125 trees below 1.5m height and 12 trees above 1.5m height but below 0.1m^3^ volume were also filtered from all analyses because the photogrammetry resolution was insufficient to reliably measure them. As a result of all exclusion steps, 49 families present in the garden were not represented in our dataset. Our final sample size was 1,170 trees from 573 families.

### 2.6. Calculating phenotypic trait values from remotely sensed data

#### 2.6.1. Canopy volume

We derived canopy volume by calculating the sum of all CHM pixel heights within the tree canopy boundaries and multiplying by the area of a pixel (7.255 x 10^-4^). To do so, we cropped and masked the CHM raster to each canopy polygon using function crop() with the “mask” option in R package terra.

#### 2.6.2. Greenness

Using the multispectral imaging data, we first calculated greenness (Normalized Vegetation Difference Index; NDVI (Gao 2000)) for each pixel within the tree canopy boundaries:

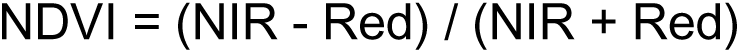

We then calculated two metrics to summarize greenness of each canopy: (1) average greenness of pixels within a canopy and (2) percent pixels with high greenness (greenness _≥_ upper quartile greenness of all pixels). Prior to calculating canopy greenness metrics, we excluded all pixels with greenness < 0.6 because any lower values are unlikely to represent living vegetation (Gao 2000). In addition, because the understory consisted of annual grasses that had died by the time of data collection, we are confident that greenness values only applied to the tree canopies. To ensure that canopy average greenness was not biased by trees with exposed bare branches or dead leaves, we also tested the correlation between average canopy greenness and percentage of the canopy with greenness in the third quartile.

### 2.7. Statistical analysis

All statistical analyses were run using R version 4.5.0 (R Core Team 2025) in RStudio version 2025.05.0 (Posit team 2025). All figures were plotted using function plot_model() in R package “sjPlot” version 2.8.17 (Lüdecke 2024), which imports package ggplot2 version 4.0.0 (Wickham 2016).

#### 2.7.1. Validating drone-derived size measurements

We used linear models to compare (a) raw drone-derived height estimates and (b) modeled drone-derived height estimates with the full dataset of field-collected height values. We then compared the accuracy of the drone-derived height estimates to our previous allometric estimates of height from DBH (see Goetz et al. 2026a for details on allometric fitting). We validated size-related metrics against the ground-truth data and fit a regression model (function lm() in package “stats”) to relate drone-derived height to hand-measured height (equation: Ht_field_= ß_0_ + ß_1_ * Ht_drone_; fitted on a randomly sampled 50% of the data). Finally, we compared the drone-derived canopy volume values to the field-collected height and DBH values using GAMs (gamma distribution with log link, to ensure that all predicted values are > 0; function bam() in R package “mgcv” version 1.9-4; Wood 2011, 2017, Wood et al. 2015) to determine whether they scaled as expected.

#### 2.7.2. Testing whether short-term greenness is associated with single-season and lifetime growth

To determine whether greenness (an instantaneous measure of tree health) is associated with growth at three different time scales, we fitted linear mixed models to compare canopy greenness with canopy volume (see above for details) and relative growth rates calculated from field-measured tree height. We tested both short-term growth rate (calculated based on values at the end of the 2023 and 2024 growing seasons) and cumulative growth rate (calculated based on values at the end of the 2014 and 2024 growing seasons). As an additional cumulative growth rate measure, we also tested associations with cumulative growth between 2014 and 2017 to test whether growth differences in early years led to canopy size and greenness differences in later years. To correct for the fact that some trees were taller than others at the beginning of the study (due to maternal effects on seeds or environmental differences while growing in the greenhouse/lath house), models also accounted for the seedling height prior to planting in the common garden. In addition, models accounted for planting block within the garden (fixed effect) and family (random effect). For a detailed description of data and calculations used to derive growth rates, see Goetz et al. 2026a. Finally, we tested whether families’ average volume and greenness were associated with percent survival.

#### 2.7.3. Testing for phenotypic trait differences among families

We tested whether phenotypic trait values (volume, greenness, lateral-vertical index, and canopy continuity indices) differed among families, using ANOVAs (function aov() in R package “stats” version 4.6.0; R Core Team 2025) that accounted for differences among planting blocks in the gardens. We then calculated quantitative trait differentiation (*Q_ST_*) and narrow-sense heritability (*h^2^*) for each trait by fitting linear mixed models with family as a random effect and planting block as a fixed effect and partitioning the variance components (Falconer and Mackay 2009). To correct for the fact that some trees were taller than others at the beginning of the study (due to maternal effects on seeds or differences while growing in the greenhouse/lath house), models also accounted for the seedling height prior to planting in the common garden.

#### 2.7.4. Testing for effects of origin climate on volume and greenness

We tested whether canopy volume and greenness were associated with three origin climate variables: T_max_, T_min_, and precipitation (see “Climates of origin” above for more detail) using linear models (function lm() in R package “stats”) that accounted for environmental differences among planting blocks. Because all trees within a family have the same origin climate, we did not include family as a blocking factor in the models.

## 3. RESULTS

### 3.1. Background: Remotely sensed tree height and canopy volume scaled predictably with ground measurements

Drone-derived tree height scaled closely with ground-measured tree height (Figure 3A). While drone-derived heights were consistently lower than ground-measured heights (Figure 3A; Table S1), the slope of the regression line was very close to 1, showing a lack of bias in the measurements. Furthermore, drone-estimated heights (Figure 3A) were more accurate and precise than our allometric estimates of height from DBH (Figure 3B) in terms of R^2^, slope, and homoscedasticity. Specifically, height is underestimated in small trees and overestimated in large trees, with variance increasing with height. (Refer to Figure S2 for the correlation matrix comparing all derived metrics.)

**Figure 3:**
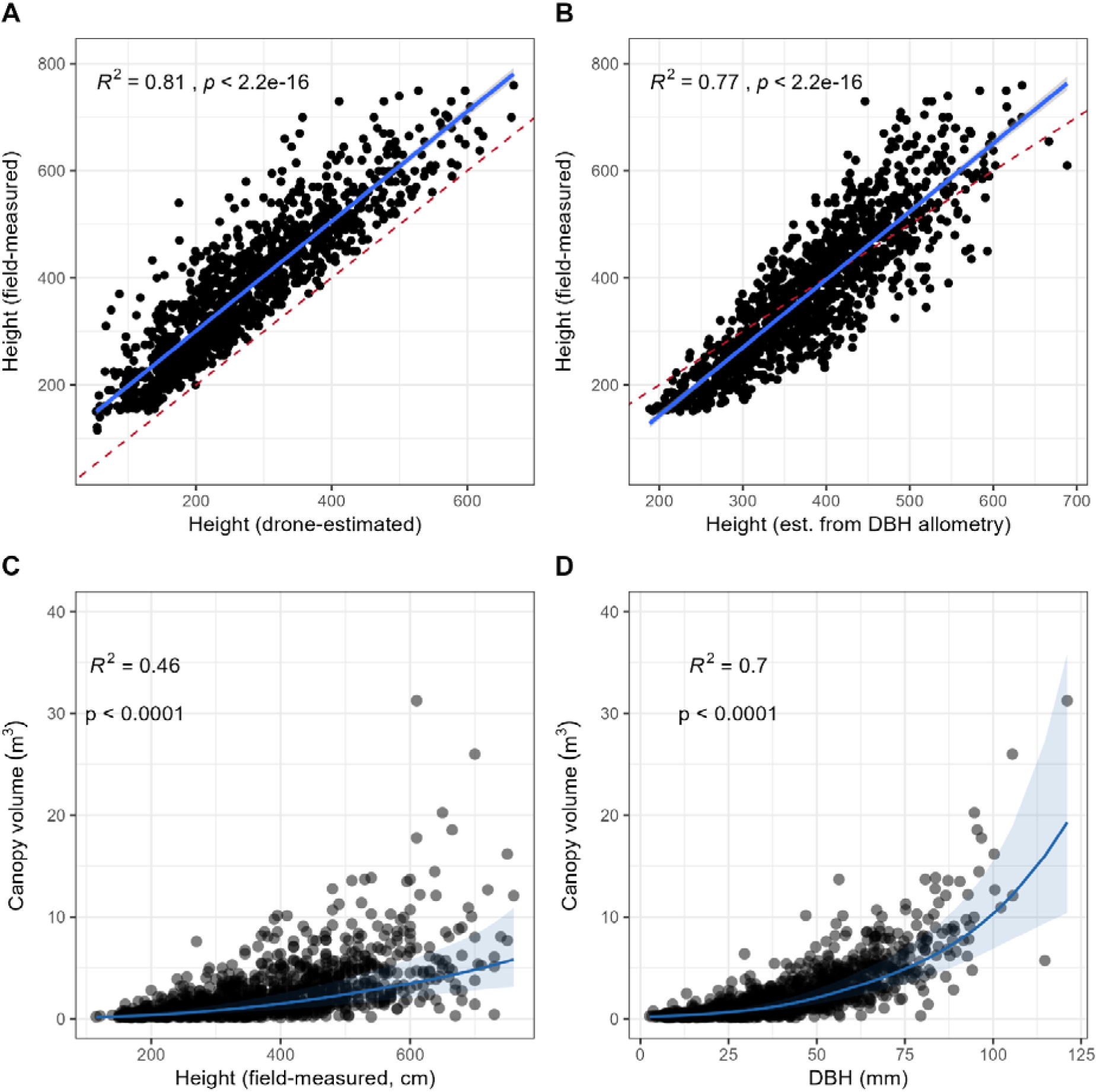
Tree size metrics from aerial photogrammetry scale as predicted with ground measurements. **(A-B)** Validation of drone-derived height estimates. Each point represents one tree in the garden; solid blue line + shading indicates regression line and 95% confidence interval. Dashed red line shows the 1:1 line. All measurements are in cm. **(A)** Raw drone-derived heights scale very closely with field-measured heights (ß_1_ = 0.999), but drone-derived heights are consistently ∼50cm lower than field-measured heights. **(B)** Height estimated from DBH using an allometric equation (Goetz et al. 2026) shows lower concordance with the field-measured heights than does the drone-derived height metric. **(C-D)** Scaling relationships between field-measured height **(C)** and drone-derived canopy volume and field-measured diameter at breast height (DBH) and drone-derived canopy volume **(D)**, estimated via generalized additive model (gamma distribution with log link) accounting for garden planting block. Both relationships are statistically significant, but DBH scales more closely than height with canopy volume and exhibits less heteroscedasticity as tree height increases.

Field-measured height and DBH also scaled predictably with drone-derived canopy volume and were statistically significant when modeled via GAMs that accounted for the effect of planting block (Figure 3C, 3D). Both fitted models were roughly exponential, but the height/volume relationship breaks down as tree height increases while DBH continues to closely predict volume in larger trees (the DBH model explains 23% more of the data variance than the height model). This discrepancy is expected because trees of similar height may vary drastically in volume due to differences in lateral growth; adult *Q. lobata* trees tend to have wide canopies relative to their height (Pavlik et al. 1991). Details of models relating canopy volume to height and DBH are provided in Tables S2 and S3.

### 3.2. Greenness is associated with growth across multiple temporal scales

Greenness was also significantly associated with canopy volume and both short-term (2023-2024) and cumulative (2014-2017; 2014-2024) growth rates when accounting for initial tree height of each growth period, planting block, and family (Figure 4). Average canopy greenness and percentage of the canopy with high greenness were strongly correlated and showed very similar relationships with growth variables, so we conclude that canopy greenness accurately represents the health of leaves instead of being confounded by bare spots in the canopy (Figure S3). See Tables S4-S7 for model summaries.

**Figure 4.**
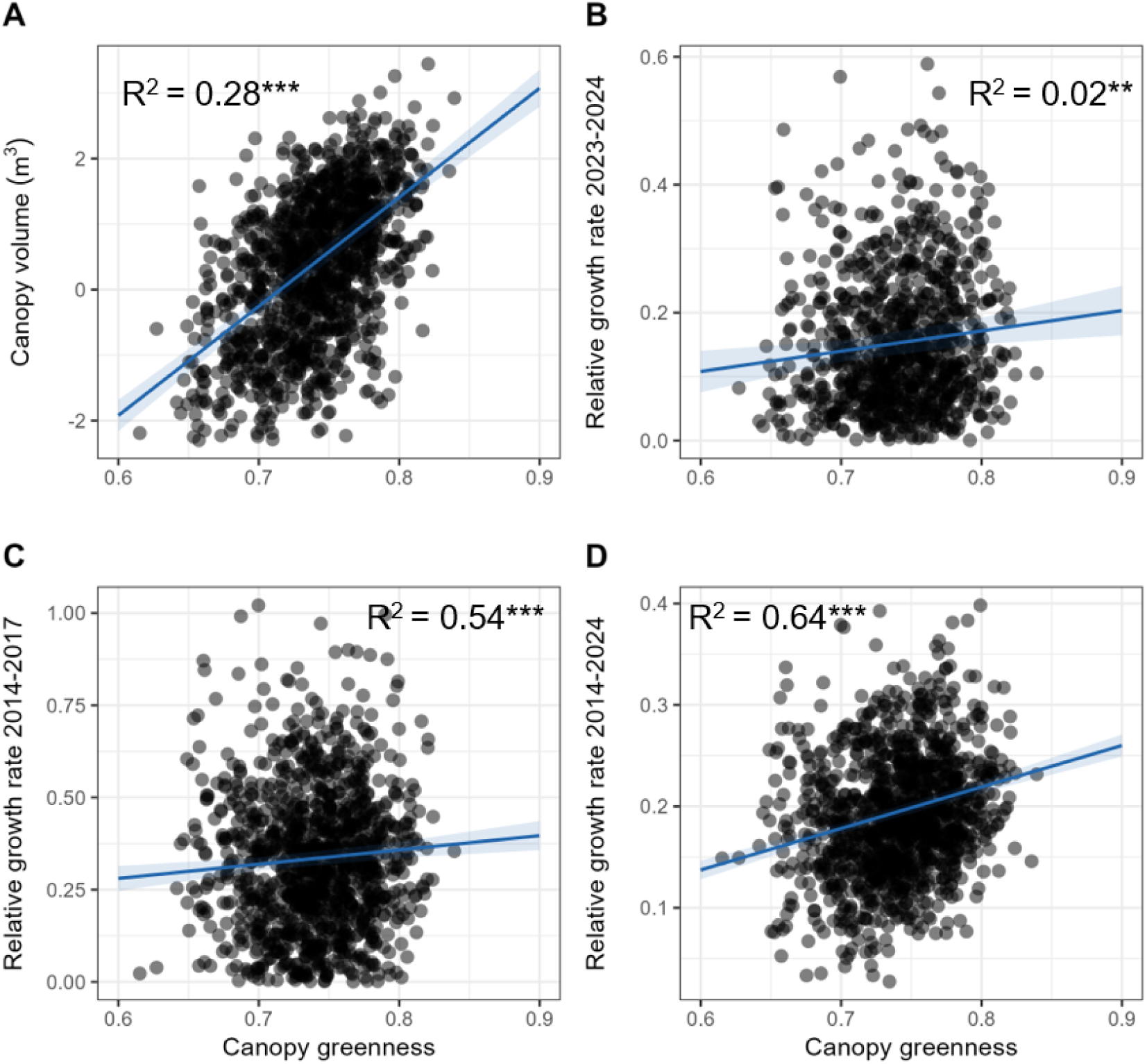
Canopy greenness is associated with canopy volume, recent relative growth rate, and lifetime relative growth rate. Relationships between canopy greenness and **(A)** canopy volume; and **(B-D)** relative growth rates. Relative growth rates were calculated from height or estimated height in three time periods: prior growing season (2023-2024), early cumulative (2014-2017), and lifetime-to-date cumulative (2014-2024). Marginal effect of each predictor variable of each linear mixed model is shown; all models also included the following terms: tree height at beginning of growth period, planting block, and family (random effect). All relationships are statistically significant (p < 0.05); best-fit lines and 95% confidence intervals are shown.

### 3.3. Phenotypes differed by family

Canopy volume, height, and greenness were significantly differentiated by family (Table 1). Differences among families explained much of the variance in traits, with *Q_ST_* _≥_ 0.60 for all traits, and *h*^2^ was particularly high for greenness (86%)(Table 2). Thus, we find support that families differ across the species range and that these polygenic traits are heritable, making them appropriate to quantify in the context of selection.

**Table 1.** ANOVAs testing differences in remotely sensed phenotypic traits among half-sib families (modeled as a random effect) of *Q. lobata* growing in a common garden. Models accounted for environmental differences among planting blocks (fixed effect). The fixed effect of tree height prior to outplanting was also included in the model to account for maternal effects on trees’ early growth.

|  |  | *df* | *SS* | *MS* | *F* | *p* |
| --- | --- | --- | --- | --- | --- | --- |
| **Log(Canopy volume)** | Planting block | 4 | 9.45 | 2.36 | 2.24 | 0.06 |
|  | Family | 572 | 841.39 | 1.47 | 1.39 | < 0.001 |
|  | Initial height | 1 | 5.54 | 5.54 | 5.25 | 0.02 |
|  | Residuals | 590 | 622.76 | 1.06 |  |  |
| **Height** | Planting block | 4 | 19.73 | 4.93 | 4.23 | < 0.001 |
|  | Family | 572 | 857.72 | 1.50 | 1.28 | < 0.001 |
|  | Initial height | 1 | 1.73 | 1.73 | 1.48 | 0.22 |
|  | Residuals | 590 | 688.63 | 1.17 |  |  |
| **Greenness** | Planting block | 4 | 0.11 | 0.03 | 29.22 | < 0.001 |
|  | Family | 572 | 0.89 | < 0.001 | 1.66 | < 0.001 |
|  | Initial height | 1 | < 0.001 | < 0.001 | 0.34 | 0.56 |
|  | Residuals | 590 | 0.56 | < 0.001 |  |  |

**Table 2.** Variance components and derived quantitative trait differentiation (*Q_ST_*) and narrow-sense heritability (*h^2^*) values of remotely sensed phenotypic traits measured among *Q. lobata* half-sib families in a common garden, estimated from a linear mixed model testing the random effects of family and planting block. The fixed effect of tree height prior to outplanting was also included in the model to account for maternal effects on trees’ early growth. V_A_: Additive genetic variance (4*V_Family_). V_P_: Total phenotypic variance (V_Family_ + V_Planting block_ + V_Residual_). Q_ST_: V_P_ / (V_P_ + 2*V_A_).

| | $V_A$ | $V_P$ | $Q_{ST}$ | $h^2$ |
| --- | --- | --- | --- | --- |
| **Log(Canopy volume)** | 0.84 | 1.25 | 0.43 | 67% |
| **Greenness** | 0.0012 | 0.0014 | 0.37 | 86% |
| **Height** | 0.67 | 1.33 | 0.50 | 50% |

### 3.4. Canopy phenotypes were associated with origin climates

Canopy volume and greenness were both significantly associated with summer maximum temperature, average winter temperature, and total annual precipitation of the origin site (Figure 5). Trees from climates with hotter summers (Figure 5A) and less precipitation (Figure 5C) had greater volume than trees from cooler and wetter climates, while the greatest volume was observed in trees from sites with similar winter temperature to the garden environment (Figure 5B). Trees with the greatest volume originated from hotter and drier climates than the garden. In contrast, trees originating from sites with similar winter temperatures as the garden had the highest volume.

**Figure 5.**
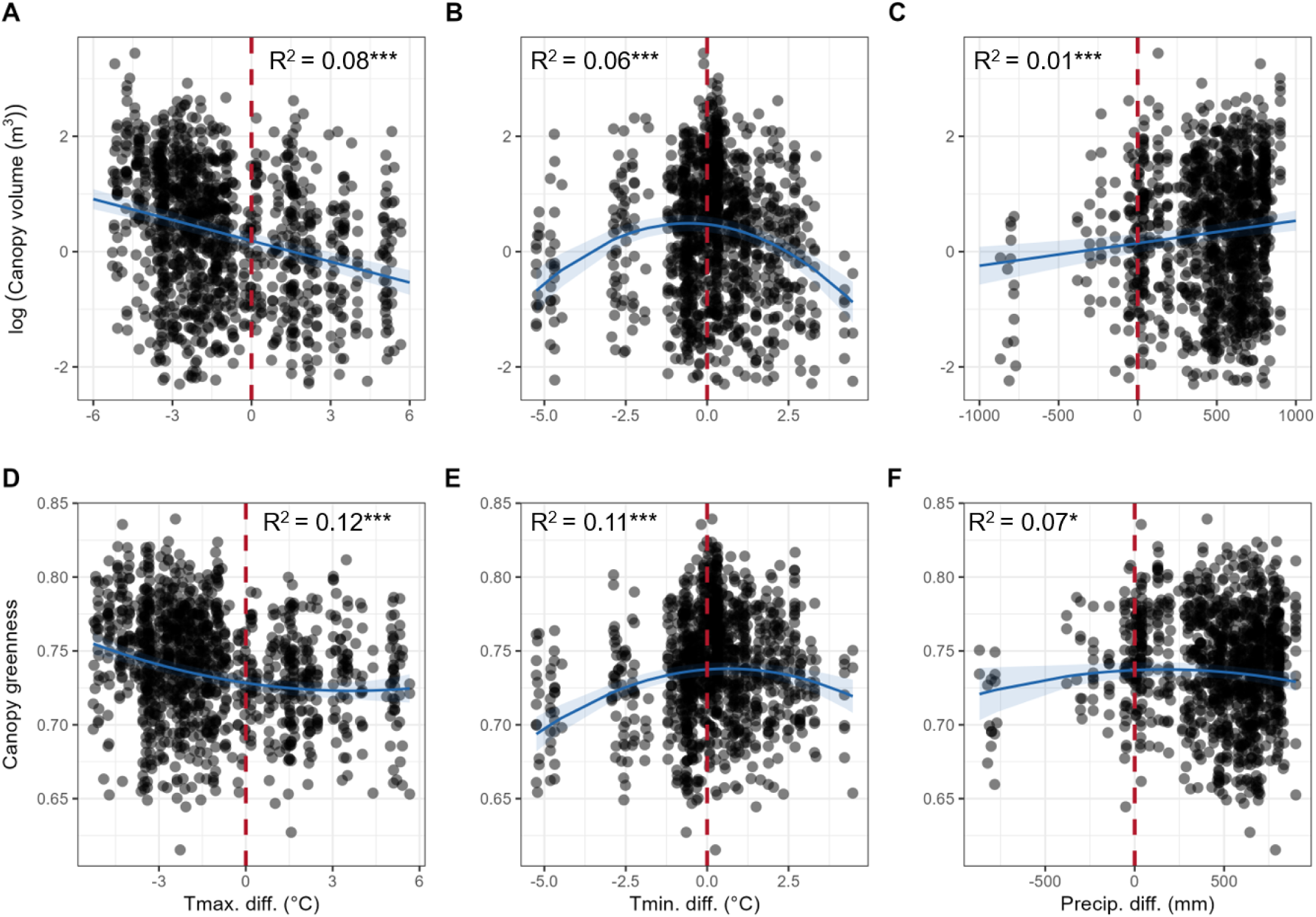
Canopy volume and greenness are associated with transfer distance between origin climate and garden conditions (2014-2024 average). (**A, D)** summer maximum temperature. (**B, E)** winter mean temperature. (**C, F)** cumulative annual precipitation. Blue curve represents the effect of climate on canopy volume **(A-C)** and greenness **(D-F)** calculated from linear models. Vertical dotted red lines represent zero transfer distance, i.e., garden climate matches source climate. Curved lines denote a significant quadratic term, while straight lines denote a significant linear term only. Shaded intervals are 95% confidence intervals. Models also accounted for planting block.

Greenness/climate relationships were very similar to volume/climate relationships, with consistent high greenness among trees from origins with hotter summers (Figure 5D) and peak greenness at intermediate values of winter mean temperature (Figure 5E). Unlike canopy volume, the highest greenness was observed in trees from origins with intermediate precipitation (Figure 5F). The effects of temperature transfer distances on both volume and greenness were stronger than the effect of precipitation. Model summaries are provided in Tables S8-S13.

## 4. DISCUSSION

The 12-year-old *Q. lobata* trees originating from climates with hot summers, moderate winters, and low annual precipitation showed the highest incremental and cumulative performance in a common garden. We obtained this evidence using high-throughput phenotypic measurements of canopy volume derived from drone imagery, which exhibited significant genetic differences associated with maternal climate. In addition, trends in greenness suggest that trees originated from warmer climates had greater vigor and/or less stress than those from cooler climates. Collectively, these findings indicate that *Q. lobata* is likely to become increasingly mismatched to future climates.

### 4.1. Spectral measurements captured functional phenotypes linked to variation in long- and short-term tree performance

The association between canopy greenness and volume suggests a meaningful link between functional performance and aboveground biomass in *Q. lobata*. Greenness, a measure of vigor and/or stress in leaves linked with photosynthetic activity, represents short-term functional performance (Sellers 1985, Kothari and Schweiger 2022). The finding that greenness is associated with canopy volume suggests that greenness may reflect variation in relative growth rate that leads to variation in canopy volume. Thus, the association between greenness and both short- and long-term growth rates indicates that higher individual photosynthesis may result in higher growth per year, which in turn is predictive of faster long-term growth (Wang et al. 2004). Early differentiation in growth predicting long-term differentiation in growth is also consistent with our previous finding that early growth was highly predictive of total growth to date (Goetz et al. 2026a). Thus, this functional metric of short-term leaf function at the whole- canopy level provides useful information about cumulative growth and tree response to the garden environment.

### 4.2. Canopy volume and greenness were genetically differentiated and heritable

The drone images of aboveground physical size accurately detected previously identified genetic differentiation in growth in the common garden, measured by changes in height and DBH as trees aged (Browne et al. 2019, Goetz et al. 2026a) and correlated with total above- and belowground biomass detected in destructively sampled trees (Ochoa et al. *in review*). Thus, we present new evidence that a canopy-scale structural trait strongly associated with lifetime fitness (Younginger et al. 2017) reflects meaningful differences in performance among *Q. lobata* families.

Genetic differentiation in greenness suggests that an additional canopy-scale phenotypic trait demonstrates a physiological dimension of performance variation that is directionally consistent with variation in size. Differences among families indicate that key physiological processes related to photosynthesis (Knipling 1970, Peñuelas and Filella 1998, Kothari and Schweiger 2022) are genetically determined and not solely a function of the environment, as has been noted in several other tree species (e.g., Ahrens et al. 2020, Corbin et al. 2025). Because we found that greenness also scaled with metrics of size, our findings suggest that genetic variation in functional performance may contribute to differences in growth and thus ultimately fitness, consistent with findings from other species (e.g., Wang et al. 2004).

### 4.3. Canopy volume and greenness were highest in trees from hot, dry origins

Because the garden’s climate conditions are close to the current-day median value across the species range (Delfino Mix et al. 2015), the fact that the best-performing trees are from hot, dry climates provides new evidence suggesting species-level maladaptation to current climates (Browne et al. 2019, Goetz et al. 2026a). By contrast, we observed a peak in canopy volume and greenness among trees with similar winter temperature to the garden, which provides further evidence of local adaptation to winter temperature but not summer temperature (Goetz et al. 2026a), likely because minimum temperature was a stronger selective pressure than maximum temperature during periods of glaciation (Gugger et al. 2013). Regardless, the maladaptive relationships between maximum temperature and both cumulative growth and short-term leaf health suggest a detrimental effect of rising temperatures on *Q. lobata* performance, which has also been noted in other tree species (e.g., Frank et al. 2017, Fréjaville et al. 2020, St. Clair et al. 2020, Gougherty et al. 2021). By testing relationships between climate, canopy volume, and canopy greenness, we show multiple avenues through which increasing temperatures under climate change may cause climatic mismatches in *Q. lobata* populations as they are exposed to temperatures further from their climate optima.

The effects of precipitation of origin on phenotypes in the common garden were weaker than the effects of temperature of origin. In contrast to the linear relationship between precipitation transfer distance and canopy volume, we found a small but significant quadratic effect of precipitation transfer distance on canopy greenness, suggesting a weak signal of local adaptation. Precipitation had a weaker effect and explained less variance in volume and greenness than maximum or minimum temperature. Single-year relative growth rate was also not significantly associated with precipitation (Figure S4), possibly because the study year was not a drought year in the garden (Figure S1). Thus, we find evidence that temperature is a stronger driver of short-term growth and leaf health than precipitation. This finding is concordant with previous evidence that maximum temperature of origin is a strong predictor of valley oak physiological response to drought (Mead et al. 2019). Temperature and precipitation are also strongly inversely correlated throughout the valley oak range (Kueppers et al. 2005, Schoenherr 2017). Increased drought is projected to be a major driver of valley oak recruitment limitation under climate change (Kueppers et al. 2005, McLaughlin and Zavaleta 2012) and more broadly a primary driver of future maladaptation throughout Mediterranean ecosystems (Matesanz and Valladares 2014, Sáenz-Romero et al. 2017, George et al. 2020, Martínez-Sancho et al. 2025). However, our results suggest that maximum temperature is a more informative indicator than precipitation for determining how population fitness is affected by climate, even if water limitation imposes more direct stress on the individual.

### 4.4. Further studies

All measurements in this study were taken at one point in time. Multiple measurements of canopy volume could indicate whether annual or less frequent measurements (e.g., at five or ten-year intervals) are sufficiently sensitive to detect changes in growth among families. It would also be useful to know whether spectral images of greenness could detect year-to-year responses to variation in temperature and precipitation, responses to extreme climate events such as multiyear drought, or differences in overall tree vigor (e.g., Depardieu et al. 2020, Cabon et al. 2023). Given that our measurements were taken at a single time point, we cannot address these questions. Nonetheless, the consistency revealed by these two novel measures of tree performance with findings from previous studies (e.g., Browne et al. 2019, Goetz et al. 2026a) illustrates the value of these high-throughput methods for assessing the effects of climate and environmental conditions on tree performance.

Quantifying canopy greenness variation in natural areas could provide an effective means of identifying seed sources for climate-conscious reforestation. Our previous work showed that identifying donor localities climatically matched to future conditions at the planting site, and selecting high-performing trees as seed sources, should result in improved population fitness (Browne et al. 2019, Goetz et al. 2026a). We suggest that potential seed sources in the wild could be compared using canopy greenness measured using drones or satellite imagery as a screening tool to identify potentially high-performing individuals. Although differences in canopy greenness among wild individuals reflect environmental effects as well as genetic differences in performance, greenness could provide a rapid indicator of tree condition under prevailing environmental conditions. This approach could help identify promising seed sources and reduce the risk of accidentally planting seeds that are maladapted to their current or future climate conditions.

### 4.4. Implications for management

Rising temperatures are likely to exacerbate climatic mismatch in valley oak trees, so identifying individuals best adapted to future conditions is of urgent concern. Consistent with previous findings (Browne et al. 2019, Goetz et al. 2026a, 2026b), these drone-produced traits suggest that *Q. lobata* individuals growing in cooler, wetter locations are at risk of climate mismatch. In addition, high individual variation in performance among lineages from similar climates demonstrates that within an environment, some trees may be better adapted than others (Mahony et al. 2020, Lind and Lotterhos 2024). As a result, it may be beneficial to incorporate phenotypic information into management practices (Vander Mijnsbrugge et al. 2010, Thomas et al. 2014). Rapid, high-throughput phenotyping of common garden studies can therefore improve our ability to empirically inform management strategies that account for past and future changes in climate.

The photogrammetry methods developed for mature coniferous forests (Young et al. 2022) can be extended for phenotyping juvenile deciduous trees in common gardens. Common garden studies are invaluable resources for conservation of wild tree populations (Wheeler et al. 2015, Gellie et al. 2016, Mahony et al. 2020, Sáenz-Romero et al. 2020), but their high associated costs present an obstacle to establishing new studies or resampling existing studies (Huxman et al. 2022). While drone-based tree phenotyping is often done by collecting lidar and hyperspectral data (e.g., Gitelson et al. 2003, Cooper et al. 2019, Lombardi et al. 2022, Grubinger et al. 2023, Corbin et al. 2025), here we demonstrate that we can effectively characterize variation in tree performance using data collected from four-band multispectral and red-green-blue optical imagery, which are measured using comparatively inexpensive sensors with less intensive computational requirements for processing and analysis. Thus, by enabling the collection of high-throughput phenotypic data with labor-efficient and relatively inexpensive tools, the methods we describe here can expand the potential of common garden studies to inform conservation practice.

## 5. Conclusions

Phenotypic traits in *Quercus lobata* derived from high-resolution, high-throughput drone-based remote sensing showed significant genetic differentiation among families and associations with climate of origin. By linking structural and functional metrics with genetic differentiation and climate-of-origin effects, we demonstrate the potential for a scalable approach to assess tree performance under climate change using high-throughput phenotyping. The correspondence between canopy greenness and tree size, significant trait differentiation among families, and the associations with climate-of-origin demonstrate that drone-derived measurements capture biological patterns consistent with those identified through conventional phenotyping. Moreover, these measurements provide complementary information on tree condition beyond what can be captured by conventional size metrics such as height or stem diameter. These methods could help identify climate-adapted seed sources and improve strategies for restoring oak populations under future climate conditions.

## AI assistance

Large language model-based tools (Microsoft Copilot, Anthropic Claude, and OpenAI ChatGPT) were used during the editing process to critique the logical flow of the text and suggest wording changes to improve clarity. No original text was generated by AI tools, and the authors take full responsibility for all content in the manuscript.

## Author contributions

VLS and JWW designed the common garden with advice from Robert Westphall; ARBG, VLS, JWW designed research questions; DJY collected and processed remotely sensed data; KKS conducted ground mapping of tree locations; VLS, JWW, ARBG, and KKS collected field data; ARBG analyzed data with input from VLS and DJY. ARBG and VLS wrote the paper, and all authors contributed revisions.

## Acknowledgements

The authors acknowledge the native peoples of California as the traditional (past, present, and future) caretakers of the ecosystems from which these data were collected. We thank Annette Delfino Mix, Courtney Canning, and Lisa Crane for their extensive work in establishing and maintaining the common garden trial. We thank Marissa Ochoa and Berenice Badillo as field team leaders during the last few years, as well as many past and present Sorklab members and volunteers who collected field data. For assistance in collecting and processing the drone imagery, we thank Jackson Anderson, Oren Nardi, and Ciro Gonzalez-Zugarazo. Finally, we thank Thomas Harbin and Rebecca Cutforth for assistance with tree mapping and Dane Sheridan for data cleaning and quality control.

The findings and conclusions in this publication are those of the authors and should not be construed to represent any official USDA or US Government determination or policy. Any use of product names is for informational purposes only and does not imply endorsement by the US Government.

## Data availability

Upon publication, data and scripts supporting this study will be made publicly available on FigShare. Tree size data will be publicly available from the Open Forest Observatory data portal (openforestobservatory.org), and all raw field data will be publicly available from the common garden project’s data portal (https://sorklab2.eeb.ucla.edu/app/). Code underlying workflows for photogrammetric processing of drone imagery are available on the Open Forest Observatory website.

