## Supplementary figures and tables for "Canopy traits reveal climate-associated genetic differentiation in a California oak common garden"

Short title: Climate of origin is associated with canopy traits and tree performance

### Supplementary Figures

**Figure S1**. Variation in total annual precipitation, 2014-2024, in a common garden of 1,395 valley oak (*Quercus lobata*) trees from 601 maternal lineages in a provenance trial in Placerville, California, USA. Trees were planted into the garden in 2014. Year 2024, in which the remotely sensed data were collected, is outlined in black. The black dotted line represents the 30-year average (1991-2020) of total annual precipitation at the common garden site. All precipitation data are taken from PRISM (prism.oregonstate.edu); refer to Methods for details.

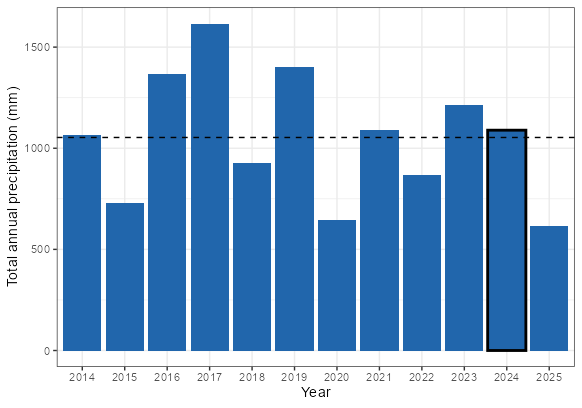

**Figure S2**. Correlation matrix of remotely sensed and field-measured size measurements on 1,395 12-year-old valley oak (*Quercus lobata*) trees from 601 maternal lineages in a provenance trial in Placerville, California, USA. “RGR” denotes relative growth rates calculated from height or estimated height at various time points (see Methods for more detail). DBH (diameter at breast height; 1.4 m) and field-measured height were collected in 2024 by field crews. Drone-estimated height, percent pixels with high NDVI (Normalized Difference Vegetation Index), NDVI, crown volume, crown width, and crown area were all measured via drone-based remote sensing in 2024. All variables were scaled as z-scores prior to testing correlations to account for differences in range among variables.

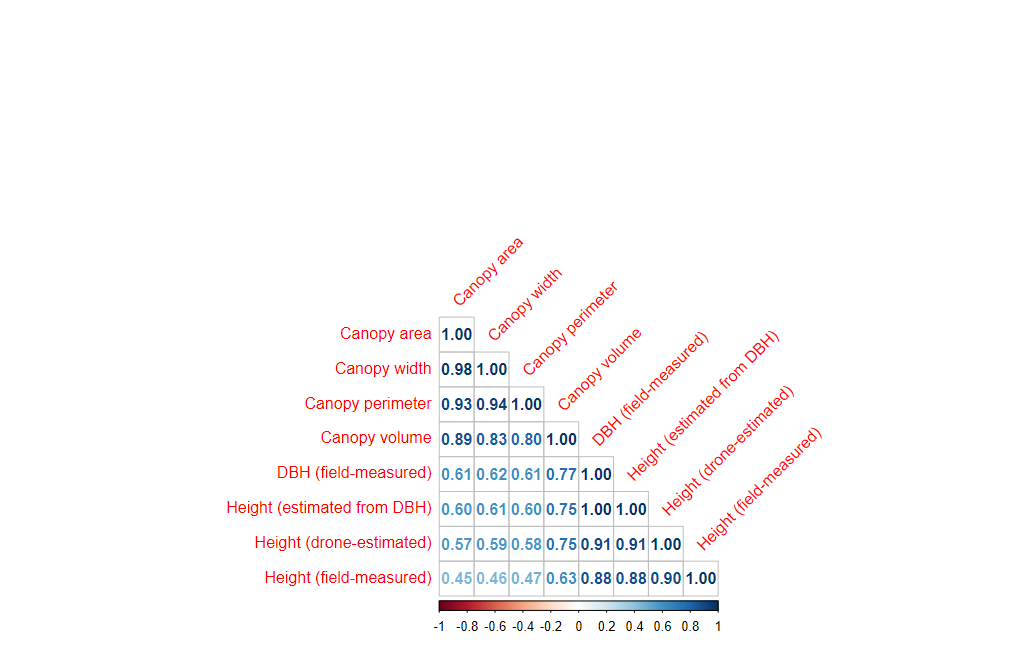

**Figure S3**. Average greenness of the entire canopy is correlated with percentage of the canopy with high greenness (normalized difference vegetation index, NDVI). (**A**) percentage of pixels above median greenness. {**B**} percentage of pixels above third-quartile greenness.

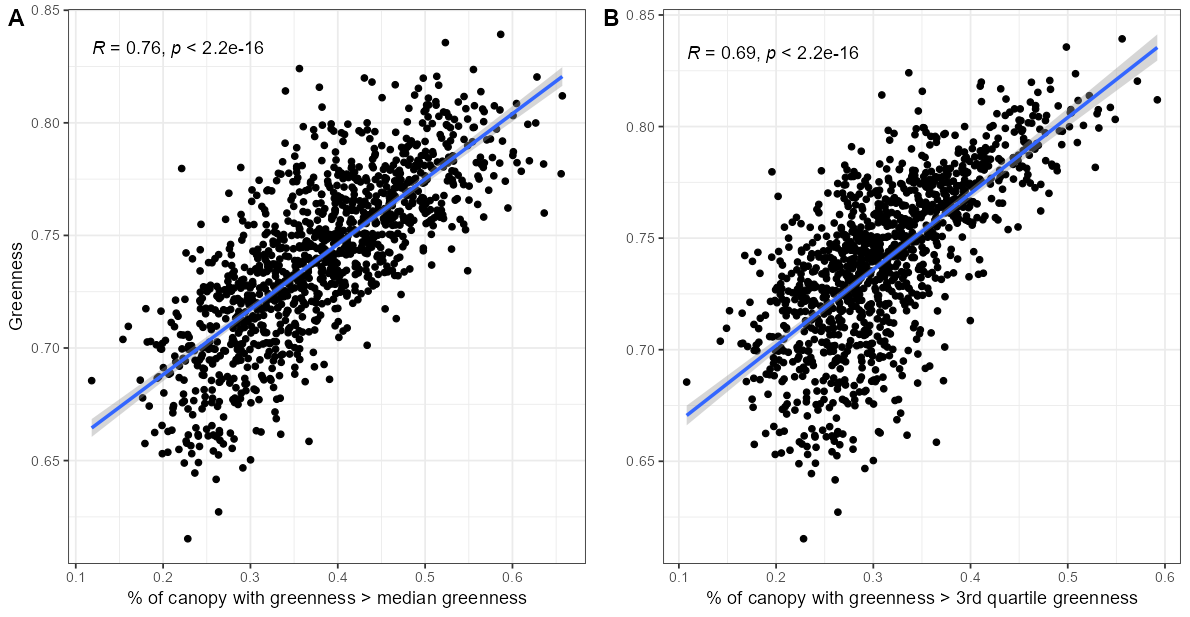

### Supplementary Tables

**Table S1**. Linear model used to adjust for consistent discrepancy between field-measured and drone-derived heights of 12-year-old valley oak trees growing in a common garden (Placerville, California, USA). We fit a regression model (function lm() in R package “stats”) to relate drone-derived height to hand-measured height (equation: Ht_field_= ß_0_ + ß_1_ * Ht_drone_; fitted on a randomly sampled 50% of the data).

|  | **Height (drone-derived; cm)** | | |
| --- | --- | --- | --- |
| *Predictors* | *Estimates* | *CI* | *p* |
| Height (field-measured; cm) | 0.79 | 0.76 – 0.83 | **<0.001** |
| (Intercept) | -18.32 | -32.59 – -4.05 | **0.012** |
| Observations | 547 | | |
| R^2^ / R^2^ adjusted | 0.793 / 0.792 | | |

**Table S2**. Details of generalized additive model testing relationship between field-measured tree height and drone-derived canopy volume. “Block” denotes planting block within the garden and was included to account for unintended environmental differences among sections of the garden. Family (maternal lineage) was added as a random effect.

|  | **Canopy volume** | | |
| --- | --- | --- | --- |
| *Predictors* | *Estimates* | *CI* | *p* |
| Smooth term (Height cm) |  |  | **<0.001** |
| Smooth term (Family) |  |  | **<0.001** |
| (Intercept) | 1.69 | 1.56 – 1.83 | **<0.001** |
| Block [2] | 1.12 | 1.00 – 1.25 | 0.052 |
| Block [3] | 1.14 | 1.01 – 1.28 | **0.030** |
| Block [4] | 1.17 | 1.05 – 1.31 | **0.004** |
| Block [5] | 1.04 | 0.93 – 1.16 | 0.540 |
| Observations | 1395 | | |
| R^2^ | 0.530 | | |

**Table S3**. Details of generalized additive model testing relationship between field-measured diameter at breast height (1.4m; DBH) and drone-derived canopy volume. “Block” denotes planting block within the garden, and was included to account for unintended environmental differences among sections of the garden. Family (maternal lineage) was added as a random effect.

|  | **Canopy volume** | | |
| --- | --- | --- | --- |
| *Predictors* | *Estimates* | *CI* | *p* |
| Smooth term (DBH) |  |  | **<0.001** |
| Smooth term (Family; random effect) |  |  | **0.016** |
| (Intercept) | 1.81 | 1.70 – 1.94 | **<0.001** |
| Block [2] | 1.09 | 0.99 – 1.21 | 0.075 |
| Block [3] | 1.08 | 0.98 – 1.20 | 0.133 |
| Block [4] | 1.12 | 1.02 – 1.24 | **0.018** |
| Block [5] | 1.06 | 0.96 – 1.18 | 0.227 |
| Observations | 1319 | | |
| R^2^ | 0.707 | | |

**Table S4**. Full model summaries of the effects of canopy volume, tree height, planting block, and maternal family on average canopy greenness (NDVI) and percent of canopy with high greenness (highest quartile NDVI).

|  | **NDVI** | | | **Percent canopy with highest-quartile NDVI** | | |
| --- | --- | --- | --- | --- | --- | --- |
| *Predictors* | *Estimates* | *CI* | *p* | *Estimates* | *CI* | *p* |
| Smooth term (vol) |  |  | **<0.001** |  |  | **<0.001** |
| Smooth term (Height cm) |  |  | **<0.001** |  |  | **<0.001** |
| Smooth term (Family) |  |  | **<0.001** |  |  | **<0.001** |
| (Intercept) | 0.72 | 0.72 – 0.72 | **<0.001** | 0.25 | 0.24 – 0.26 | **<0.001** |
| Block [2] | 1.00 | 0.99 – 1.00 | 0.055 | 0.97 | 0.92 – 1.02 | 0.217 |
| Block [3] | 1.00 | 1.00 – 1.01 | 0.764 | 1.01 | 0.96 – 1.06 | 0.696 |
| Block [4] | 1.01 | 1.01 – 1.02 | **<0.001** | 1.10 | 1.05 – 1.15 | **<0.001** |
| Block [5] | 1.03 | 1.02 – 1.03 | **<0.001** | 1.26 | 1.20 – 1.32 | **<0.001** |
| Observations | 1395 | | | 1395 | | |
| R^2^ | 0.592 | | | 0.409 | | |

**Table S5.** Full model summaries of the effects of tree relative growth rate (2023-2024 growing season), initial tree height, planting block, and maternal family on average canopy greenness (NDVI) and percent of canopy with high greenness (highest quartile NDVI).

|  | **NDVI** | | | **Percent canopy with highest-quartile NDVI** | | |
| --- | --- | --- | --- | --- | --- | --- |
| *Predictors* | *Estimates* | *CI* | *p* | *Estimates* | *CI* | *p* |
| Smooth term (RGR_2023-2024_) |  |  | **<0.001** |  |  | **0.008** |
| Smooth term (2023 ht) |  |  | **<0.001** |  |  | **<0.001** |
| Smooth term (Family) |  |  | **<0.001** |  |  | **<0.001** |
| (Intercept) | 0.72 | 0.72 – 0.73 | **<0.001** | 0.25 | 0.24 – 0.26 | **<0.001** |
| Block [2] | 0.99 | 0.99 – 1.00 | 0.061 | 0.96 | 0.91 – 1.01 | 0.142 |
| Block [3] | 1.00 | 1.00 – 1.01 | 0.143 | 1.02 | 0.96 – 1.09 | 0.435 |
| Block [4] | 1.02 | 1.01 – 1.02 | **<0.001** | 1.11 | 1.06 – 1.17 | **<0.001** |
| Block [5] | 1.03 | 1.03 – 1.04 | **<0.001** | 1.28 | 1.21 – 1.35 | **<0.001** |
| Observations | 1052 | | | 1052 | | |
| R^2^ | 0.558 | | | 0.443 | | |

**Table S6**. Full model summaries of the effects of tree relative growth rate (early cumulative growth, 2014-2017), tree height, planting block, and maternal family on average canopy greenness (NDVI) and percent of canopy with high greenness (highest quartile NDVI).

|  | **NDVI** | | | **Percent canopy with highest-quartile NDVI** | | |
| --- | --- | --- | --- | --- | --- | --- |
| *Predictors* | *Estimates* | *CI* | *p* | *Estimates* | *CI* | *p* |
| Smooth term (RGR_2014-2017_) |  |  | **<0.001** |  |  | **<0.001** |
| Smooth term (2014 ht) |  |  | **<0.001** |  |  | **<0.001** |
| Smooth term (Family) |  |  | **<0.001** |  |  | **<0.001** |
| (Intercept) | 0.72 | 0.72 – 0.73 | **<0.001** | 0.26 | 0.25 – 0.27 | **<0.001** |
| Block [2] | 0.99 | 0.99 – 1.00 | **0.028** | 0.95 | 0.90 – 1.00 | 0.052 |
| Block [3] | 1.00 | 0.99 – 1.00 | 0.479 | 0.97 | 0.91 – 1.03 | 0.321 |
| Block [4] | 1.02 | 1.01 – 1.02 | **<0.001** | 1.11 | 1.05 – 1.16 | **<0.001** |
| Block [5] | 1.03 | 1.02 – 1.03 | **<0.001** | 1.23 | 1.17 – 1.29 | **<0.001** |
| Observations | 1391 | | | 1391 | | |
| R^2^ | 0.407 | | | 0.286 | | |

**Table S7**. Full model summaries of the effects of tree relative growth rate (total cumulative growth, 2014-2024), tree height, planting block, and maternal family on average canopy greenness (NDVI) and percent of canopy with high greenness (highest quartile NDVI).

|  | **NDVI** | | | **Percent canopy with highest-quartile NDVI** | | |
| --- | --- | --- | --- | --- | --- | --- |
| *Predictors* | *Estimates* | *CI* | *p* | *Estimates* | *CI* | *p* |
| Smooth term (RGR_2014-2024_) |  |  | **<0.001** |  |  | **<0.001** |
| Smooth term (2014 ht) |  |  | **<0.001** |  |  | **<0.001** |
| Smooth term (Family) |  |  | **<0.001** |  |  | **<0.001** |
| (Intercept) | 0.72 | 0.72 – 0.72 | **<0.001** | 0.25 | 0.24 – 0.26 | **<0.001** |
| Block [2] | 1.00 | 0.99 – 1.00 | 0.170 | 0.96 | 0.91 – 1.01 | 0.137 |
| Block [3] | 1.00 | 0.99 – 1.01 | 0.937 | 0.99 | 0.93 – 1.04 | 0.619 |
| Block [4] | 1.02 | 1.01 – 1.02 | **<0.001** | 1.11 | 1.06 – 1.16 | **<0.001** |
| Block [5] | 1.03 | 1.02 – 1.04 | **<0.001** | 1.26 | 1.20 – 1.33 | **<0.001** |
| Observations | 1393 | | | 1393 | | |
| R^2^ | 0.524 | | | 0.393 | | |

**Table S8**. Full model (generalized additive model; GAM) summary of the effects of summer maximum temperature of locations of 601 maternal valley oak (*Quercus lobata*) trees on log-transformed canopy volume of 12-year-old progeny growing in a common garden (Placerville, California, USA), accounting for differences among planting blocks and maternal families. Canopy volume was measured via drone in 2024; temperature of the maternal site (1951-1980 average) and other environmental variables reported in this study were taken from the Basin Characterization Model (Flint et al. 2013). See Methods for more details.

|  | **Log(canopy volume)** | | |
| --- | --- | --- | --- |
| *Predictors* | *Estimates* | *CI* | *p* |
| Smooth term (summer max. temp.) |  |  | **<0.001** |
| Smooth term (Family) |  |  | **<0.001** |
| (Intercept) | 0.44 | 0.31 – 0.56 | **<0.001** |
| Block [2] | 0.02 | -0.15 – 0.20 | 0.787 |
| Block [3] | -0.19 | -0.37 – -0.00 | **0.047** |
| Block [4] | 0.11 | -0.06 – 0.28 | 0.207 |
| Block [5] | -0.16 | -0.34 – 0.02 | 0.080 |
| Observations | 1395 | | |
| R^2^ | 0.235 | | |

**Table S9**. Full model summary of the effects of mean winter temperature of the maternal site on log-transformed canopy volume, accounting for differences among planting blocks and maternal families.

|  | **Log(canopy volume)** | | |
| --- | --- | --- | --- |
| *Predictors* | *Estimates* | *CI* | *p* |
| Smooth term (winter mean temp.) |  |  | **<0.001** |
| Smooth term (Family) |  |  | **<0.001** |
| (Intercept) | 0.45 | 0.33 – 0.57 | **<0.001** |
| Block [2] | 0.01 | -0.16 – 0.19 | 0.873 |
| Block [3] | -0.20 | -0.38 – -0.01 | **0.037** |
| Block [4] | 0.12 | -0.06 – 0.29 | 0.184 |
| Block [5] | -0.20 | -0.38 – -0.03 | **0.024** |
| Observations | 1395 | | |
| R^2^ | 0.238 | | |

**Table S10**. Full model summary of the effects of annual precipitation at the maternal site on log-transformed canopy volume, accounting for differences among planting blocks and maternal families.

|  | **Log(canopy volume)** | | |
| --- | --- | --- | --- |
| *Predictors* | *Estimates* | *CI* | *p* |
| Smooth term (precipitation) |  |  | **<0.001** |
| Smooth term (Family) |  |  | **<0.001** |
| (Intercept) | 0.44 | 0.31 – 0.57 | **<0.001** |
| Block [2] | 0.01 | -0.17 – 0.19 | 0.879 |
| Block [3] | -0.19 | -0.37 – 0.00 | 0.052 |
| Block [4] | 0.11 | -0.07 – 0.28 | 0.219 |
| Block [5] | -0.18 | -0.36 – -0.00 | **0.049** |
| Observations | 1395 | | |
| R^2^ | 0.238 | | |

**Table S11**. Full model summary of the effects of summer maximum temperature of the maternal site on average canopy greenness, accounting for differences in canopy volume and among planting blocks and maternal families.

|  | **NDVI** | | |
| --- | --- | --- | --- |
| *Predictors* | *Estimates* | *CI* | *p* |
| Smooth term (summer max. temp.) |  |  | **<0.001** |
| Smooth term (Family) |  |  | **<0.001** |
| Log(Canopy volume) | 0.02 | 0.91 – 1.01 | 0.137 |
| (Intercept) | 0.72 | 0.24 – 0.26 | **<0.001** |
| Block [2] | -0.00 | 0.93 – 1.04 | 0.619 |
| Block [3] | -0.00 | 1.06 – 1.16 | **<0.001** |
| Block [4] | 0.01 | 1.20 – 1.33 | **<0.001** |
| Block [5] | 0.02 |  | **<0.001** |
| Observations | 1395 | | |
| R^2^ | 0.555 | | |

**Table S12**. Full model summary of the effects of mean winter temperature (winter mean temp.) of the maternal site on average canopy greenness, accounting for differences in canopy volume and among planting blocks and maternal families.

|  | **NDVI** | | |
| --- | --- | --- | --- |
| *Predictors* | *Estimates* | *CI* | *p* |
| Smooth term (winter mean temp.) |  |  | **<0.001** |
| Smooth term (Family) |  |  | **<0.001** |
| Log(Canopy volume) | 0.02 | 0.91 – 1.01 | 0.137 |
| (Intercept) | 0.72 | 0.24 – 0.26 | **<0.001** |
| Block [2] | -0.00 | 0.93 – 1.04 | 0.619 |
| Block [3] | -0.00 | 1.06 – 1.16 | **<0.001** |
| Block [4] | 0.01 | 1.20 – 1.33 | **<0.001** |
| Block [5] | 0.02 |  | **<0.001** |
| Observations | 1395 | | |
| R^2^ | 0.557 | | |

**Table S13**. Full model summary of the effects of annual precipitation at the maternal site on average canopy greenness, accounting for differences in canopy volume and among planting blocks and maternal families.

|  | **NDVI** | | |
| --- | --- | --- | --- |
| *Predictors* | *Estimates* | *CI* | *p* |
| Smooth term (ppt) |  |  | **<0.001** |
| Smooth term (Family) |  |  | **<0.001** |
| Log(Canopy volume) | 0.02 | 0.02 – 0.02 | **<0.001** |
| (Intercept) | 0.72 | 0.71 – 0.72 | **<0.001** |
| Block [2] | -0.00 | -0.01 – -0.00 | **0.050** |
| Block [3] | -0.00 | -0.00 – 0.00 | 0.941 |
| Block [4] | 0.01 | 0.01 – 0.01 | **<0.001** |
| Block [5] | 0.02 | 0.02 – 0.02 | **<0.001** |
| Observations | 1395 | | |
| R^2^ | 0.556 | | |
